# ClustAssess: tools for assessing the robustness of single-cell clustering

**DOI:** 10.1101/2022.01.31.478592

**Authors:** Arash Shahsavari, Andi Munteanu, Irina Mohorianu

**Author notes:** Corresponding author: I Mohorianu.

## Abstract

The transition from bulk to single-cell analyses refocused the computational challenges for high-throughput sequencing data-processing. The core of single-cell pipelines is partitioning cells and assigning cell-identities; extensive consequences derive from this step; generating robust and reproducible outputs is essential. From benchmarking established single-cell pipelines, we observed that clustering results critically depend on algorithmic choices (e.g. method, parameters) and technical details (e.g. random seeds).

We present `ClustAssess`, a suite of tools for quantifying clustering robustness both within and across methods. The tools provide fine-grained information enabling (a) the detection of optimal number of clusters, (b) identification of regions of similarity (and divergence) across methods, (c) a data driven assessment of optimal parameter ranges. The aim is to assist practitioners in evaluating the robustness of cell-identity inference based on the partitioning, and provide information for choosing robust clustering methods and parameters.

We illustrate its use on three case studies: a single-cell dataset of in-vivo hematopoietic stem and progenitors (10x Genomics scRNA-seq), in-vitro endoderm differentiation (SMART-seq), and multimodal in-vivo peripheral blood (10x RNA+ATAC). The additional checks offer novel viewpoints on clustering stability, and provide a framework for consistent decision-making on preprocessing, method choice, and parameters for clustering.

## 1 Introduction

For recent studies, single-cell sequencing took over the sequencing landscape becoming the *de facto* standard for high resolution studies. The size of resulting datasets, comprising up to millions of cells [1], offer an unprecedented level of detail, advancing the understanding of biological phenomena e.g. enabling the detailed characterisation of cell cycle [2] or zebrafish embryogenesis [3]. The technology also holds a translational promise, including for cancer patient stratification methods [4] and identification of novel therapeutic targets [5]. Major challenges for single-cell analyses remain related to data integration, scalability, and leveraging spatial information [6, 7]. Extracting information from these datasets relies on advances in computational methods, and the robustness of the subsequent outputs.

As single-cell RNA-seq (scRNA-seq) datasets sizes grew exponentially [1], clustering emerged as a central step of pipelines, either for the identification and characterisation of subpopulations (cell-types, cell-states [8], [9]), or as an intermediate step before further analyses e.g. trajectory inference [10]. Clustering is also essential to the analysis of other single-cell modalities e.g. ATAC [11], cytometry [12], lipidomics [13], as well as to single-cell multi-omics [14, 15]. Frequently used scRNA-seq clustering methods are available in pipelines like Seurat [16] and SCANPY [17], both using community detection on nearest-neighbour graphs, and Monocle [18], which provides several clustering methods, including a density-peak-based approach. Post clustering, the assignment of subpopulations to cell-types relies on cluster-specific discriminative genes, as well as known cell-type marker genes (manually curated and often validated through low-throughput methods or identified automatically using classifiers [19]).

While it is desirable to cluster cells based on biological signal, partitions can also be driven by technical nuisance effects (sequencing depth or noise [20, 21, 22]). Beyond the interplay of technical [23] and biological signals affecting clustering outputs, the results also depend on algorithmic decisions e.g. cell filtering, normalisation, feature selection, number of clusters, and the clustering algorithm [24]. Computational details, e.g. random seed, also influence the robustness of results. Currently, these decisions are often made in an *ad-hoc* manner, and vary substantially across analyses. The evaluation of a cell-partitioning, in terms of reproducibility and biological interpretation, is thus critical to ensure that relevant, functional information is captured. Ideally, clustering results would be assessed against external, ground-truth cell-type labels; however, the experimental context of most studies makes such labeling intractable. In their absence, the validity of the output relies on the robustness of clustering results.

Previously, benchmark comparisons were performed on several clustering methods, on multiple scRNA-seq data types, across feature sets [25, 26]. These studies relied on the adjusted Rand index (ARI) versus ground-truth annotations or across resulting partitions and provided valuable macro-scale information on clustering quality, stability, and speed, as well as appreciation for how acutely clustering results depend on preprocessing and the characteristics of the dataset. However, the benchmarks suggested no clustering method or pipeline as optimal across data types and conditions. Therefore, the clustering evaluation of new data, with unfamiliar characteristics, and which often lacks cell-type annotations, remains of major interest.

Here we present `ClustAssess`, a collection of tools that enable the quantification of the robustness of partitions. We focus on the evaluation of dataset-specific clustering results, in contrast to the wide-ranging and generic results provided by clustering benchmarks. We illustrate these tools on various partitions driven by choices of features, methods, on both 10x and SMART-seq scRNA-seq datasets [27, 28], as well as a multimodal single-cell RNA and ATAC dataset. We highlight several sources of variation that affect final clustering results, including method parameters, input features and approximation errors.

## 2 Materials

We illustrate the ClustAssess pipeline on the two most frequently used types of single-cell RNA-seq assays: the droplet-based 10x Chromium and the platelet-based SMART-seq i.e. (a) an in-vivo 10x Genomics dataset of hematopoiesis (available at BioStudies accession SUBS4, donor SAMEA6646090) comprising cells from spleen, peripheral blood, and blood marrow [27] and (b) an in-vitro SMART-seq dataset of human endoderm differentiation (accession ERP016000 at European Nucleotide Archive) [28]; cells from time points 0-3 of cell lines hayt, naah, vils, pahc, melw and qunz were used.

For the multimodal analysis, we illustrate the stability of partitioning results on a 10x single-cell ATAC and RNA assay from the 10x example datasets, specifically the “PBMC from a healthy donor - granulocytes removed through cell sorting (10k)” dataset (https://www.10xgenomics.com/resources/datasets/pbmc-from-a-healthy-donor-granulocytes-removed-through-cell-sorting-10-k-1-standard-1-0-0).

## 3 Methods

All analyses were performed in R v4.0.3. Packages employed throughout (including their versions) are specified in line. The analyses and benchmarking were performed on dedicated Linux servers (Debian GNU/Linux v10 and Linux kernel version 4.19.0-12-amd64).

### 3.1 Single-cell data processing and analysis

#### [10x scRNA-seq data]

The quality of raw fastq files was assessed using FastQC v0.11.3 and summarised with MultiQC v1.8 [29]; the alignment and feature quantification were performed with 10x Cell Ranger v3.1.0, using the GRCh38 v3 reference transcriptome. The distributions of sequencing depths, number of features and proportions of reads from mitochondrial (MT) genes and from ribosomal protein-coding genes (RP), per cell, were summarised in violin plots; cells with <1,000 unique features, >10% MT or <20% RP were discarded; after filtering, MT and RP genes were excluded from the count matrices. Raw expression levels were normalized with SCTransform [30]. Further quality checks included PCAs and UMAPs; raw and normalised sequencing depths, MT% and RP% were represented on a colour gradient, to assess technical variation. The Pearson residuals of the 2,000 most abundant genes were used for PCA, and the first 30 principal components (PCs) were used to calculate the UMAP.

Cells were clustered with Seurat v4.0.5 [31], Monocle v3 [32], and Monocle v2.18 [33], separately. For Seurat, a 20-shared nearest neighbour (SNN) graph was constructed on a PCA of the data, and partitioned with SLM clustering [34]. For Monocle v3, a 20-nearest neighbour graph was created on a UMAP dimensionality reduction, and clustered with Leiden algorithm [35] with resolution=5e-4. For Monocle v2, clusters were predicted with density-peak search on a tSNE dimensionality reduction, searching for the number of clusters inferred using Seurat.

#### [SMART-seq data]

The raw fastq files were evaluated analogously to 10x data. Reads were aligned to the GRCh38.p13 reference genome using STAR v2.7.6a with Ensembl v102 annotations [36]; the same annotations were used for feature quantification with featureCounts v2.0.1 [37]. The distributions of sequencing depths, number of features and MT and RP proportions, per cell, were assessed; cells with <200 unique features were discarded, genes not present in at least 3 cells were removed. Further quality checks included PCAs and UMAPs coloured by covariates e.g. donor, time point, raw and normalised sequencing depth. The normalisation was performed using SCTransform [30]; the MT and RP genes, post filtering, were excluded from the expression matrix. The analysis-specific parameters are detailed in the following sections.

#### [CellphoneDB analysis]

An SCT-normalized expression matrix was provided as input, along with Seurat and Monocle cluster assignation; the CellphoneDB CLI v2.1.7 (`method statistical_analysis`) was used for predicting interactions [38]. Interaction terms with p-value <0.05 were grouped by sender and receiver cluster, respectively, and compared across clustering outputs with JSI.

#### [Trajectory inference and pseudotime]

Slingshot v2.1.0 [39] was used to perform trajectory inference on the Mende et al. dataset [27]. Seurat and Monocle clusterings were used as input with the isolated island corresponding to Seurat cluster 15 removed; Slingshot was applied (with parameters `stretch=2, approx_points=100`) to infer cellular trajectories. The two partitions were compared with the annotations from [27] to assess the correspondence (0 as HSC/MPP-tier-1 and 3,5,9,12 as Lymphoid, Ery/Meg/Baso/MC, Myeloid, DC respectively in Seurat; 1 as HSC/MPP-tier-1 and 6,10,12,8 as Lymphoid, Ery/Meg/Baso/MC, Myeloid, DC respectively in Monocle v3; 5 as HSC/MPP-tier 1 and 1,2,4,11 as Lymphoid, Ery/Meg/Baso/MC, Myeloid, DC respectively in Monocle v2). Subsequently, HSC/MPP-tier-1 cluster was used as the start point, and each cluster corresponding to other cell types was used as end point for the slingshot trajectories. Pseudotime was calculated along the cells belonging to each branch of the resulting lineage tree. To obtain pseudotimes across the entire dataset, the median of pseudotimes was calculated, per cell, across all branches. To quantify the dependence of Slingshot pseudotime on the inferred partitioning, the resulting median pseudotime for the Seurat and the Monocle clusterings were compared using the absolute difference between the inferred pseudotimes per cell.

#### [Multimodal data processing]

A Seurat object was created on the RNA and ATAC modalities as separate assays. Any cells failing RNA/ATAC QC tests were removed; cells with 1,000<x<100,000 ATAC counts, <25,000 RNA counts, >1,000 unique genes in the RNA, <30% MT genes and <40% RP genes (thresholds selected based on inspection of distributions) were retained for downstream analysis, totalling 9,963 cells.

As for the scRNA-seq component, the MT and RP genes were removed from the counts matrix before normalisation of gene expression (using SCTransform), and calculating a PCA reduction of the RNA modality using the 2,000 most abundant genes. A 20-SNN graph was calculated using the PCA. For the ATAC modality, after normalising the data with term frequency-inverse document frequency normalisation [16], a latent semantic indexing (LSI) dimensionality reduction was calculated on the 5% most covered peaks (5,413 peaks) using Signac v1.1.0 [40]. The LSI reduction was subsequently used for the calculation of a 20-SNN graph. Finally, a multimodal weighted 20-SNN graph was calculated jointly on the PCA and LSI dimensionality reductions using the `FindMultiModalNeighbors` function in Seurat.

For each of the three generated SNN graphs, SLM community detection was employed to find clusters, with the resolution parameter set to 0.8, across 100 random seeds. Using `merge_partitions` from `ClustAssess`, partitions with ECS *>* 0.99 were merged to identify the most frequent partition, which was then used for visualisations, ECS across modalities, confusion matrices, and marker gene calculations for each modality.

### 3.2 ClustAssess pipeline

The `ClustAssess` R package comprises functions for evaluating clustering stability with regard to the number of clusters using proportion of ambiguous clusterings (Supplementary Methods S1), functions for quantifying per-observation agreement between two or more clusterings using element-centric clustering comparison (Supplementary Methods S2), as well as plotting functions for visualisation of the results. The package also provides summary assessments to evaluate the consequences of different parameters and method choices throughout the graph-based clustering pipeline, including the effects of feature set, the number of neighbours, and clustering resolution, detailed below and in the results section.

#### Dimensionality reduction

The input for the dimensionality reduction is a normalised expression matrix (methods illustrated in the manuscript and the vignette are SCTransform [30] and log-normalisation [41]). For the precise (up to machine precision) calculation of PCA the `prcomp` method was used [42] on standardised, normalised expression matrix (i.e. centred and scaled); 30 PCs were selected using the `rank` parameter. Approximate PCA was calculated using `irlba` [43] using the same number of PCs (30) as for the exact approach. The precision was set using the tolerance parameter tol (values tested: 1e-5, 1e-10 and 1e-15).

#### Graph building

The unweighted nearest neighbour graph is built using the `nn2` function (RANN package [44]). For this implementation a point is its own neighbour i.e. it is based on an actual number of *k* − 1 nearest neighbours (further implementation details on calculating neighbourhoods are presented in Supplementary Information 1); the SNN graph is built using the `FindNeighbors` function (`Seurat` package) or the `cluster_cells` function (`Monocle` package). Default parameters are used, except for `nn.method` from `Seurat`, which is set to `rann`, and weight from `Monocle`, which is set to `TRUE`.

#### Graph clustering

The clustering is assessed on the Louvain [45], Louvain with multi-level refinement [46], SLM [34] and Leiden [35] community detection methods, available as options for the `algorithm` argument in the `FindClusters` function from `Seurat`. The resolution (set with the `resolution` parameter) was evaluated for the 0.1 to 1 range. For comparisons of the quality functions, we focused on the Leiden implementation (`cluster_cells` function) in `Monocle`; the quality function was changed using the `partition_type` parameter (values used: `CPMVertexPartition, RBConfigurationVertexPartition, RBERConfigurationVertexPartition`).

Multiple steps, including the approximate PCA, non-linear dimensionality reduction, graph building and clustering, are stochastic methods; we therefore also evaluated the effect of random seeds. The solution presented as optimal in ClustAssess corresponds to the most frequent partition.

## 4 Results

### 4.1 Clustering parameters critically influence downstream results

A first case study for illustrating the analysis-induced variability is based on the Mende et al. data (10x scRNA-seq in-vivo dataset of human hematopoietic stem and progenitor cells originating from bone marrow, peripheral blood, and spleen [27]). As initial QC, the distributions of sequencing depths, number of detected features, proportions of MT and RP genes were assessed (Supplementary Table S1). Based on these distributions, cells with fewer than 1,000 features, >10% mitochondrial genes, or <20% ribosomal protein-coding genes were discarded; these filters reduced the number of cells from 15,030 to 13,189. Expression levels were subsequently normalised using SCTranform [30].

Throughout the analysis we focus on 2 pipelines: Seurat v4, Monocle v3, denoted as the Seurat pipeline and the Monocle pipeline respectively.

#### [Independent approaches for determining optimal parameters]

A recurrent, essential question in single-cell data analysis is the number of cell sub-populations. Independent of other pipeline-specific parameters, the proportion of ambiguously clustered pairs (PAC) [47], overviewed in Supplementary Methods S1, can be used to assess the robustness of partitions with regard to the number of clusters (*k*); lower PAC indicates a more stable clustering. The PAC landscape (PAC vs *k*) can be used to avoid *k* values leading to particularly high PAC (high local variability), or, when focusing on local minima, to identify a optimal range for *k*. On the Mende et al. dataset, the PAC is monotonously decreasing with *k* i.e. higher *k* results in more stable partitions (Figure S1A); however, no obvious local optimum is observed rendering this angle inconclusive.

#### [Assessment of clustering method-induced variability]

Next, we assessed the partitioning outputs obtained using standard pipelines (Monocle [32] and Seurat [31]). For the latter, applying SLM community detection, on default parameters, i.e. on the 20-nearest neighbour graph with resolution parameter 0.8, resulted in 16 clusters (Figure 4.1A). Separately, the data was clustered with Monocle v3, using Leiden clustering with `resolution=5e-4` to delineate 13 clusters. These two methods cluster certain areas of the UMAP similarly, e.g. the far right of the UMAP (cluster 3 in Seurat, cluster 6 in Monocle); however, discrepancies between the methods emerge for bridging cells (clusters 10, 12 in Seurat vs cluster 8 in Monocle) or for compact regions, for which a robust partitioning is more challenging (cluster 1 in Seurat vs clusters 3, 4, 7 in Monocle).

**Figure 4.1:**
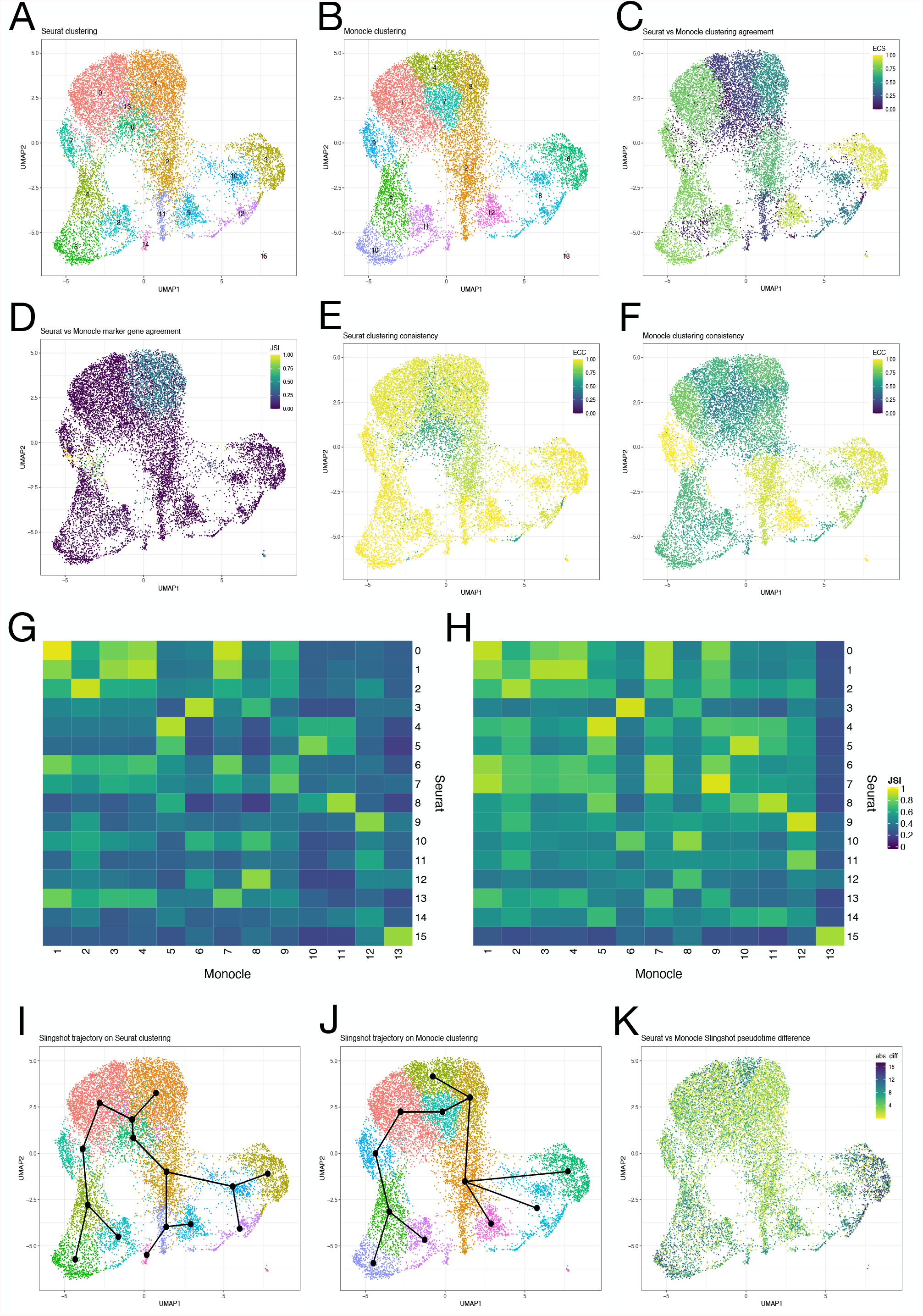
Differences between Seurat and Monocle clustering outputs, and their knock-on effects on downstream results. A-B. Seurat v4 (A) and Monocle v3 (B) clustering each cluster the Mende et al. single-cell data into 16 clusters. C. Element-centric similarity (ECS) of Seurat and Monocle clusterings quantifies per-cell clustering agreement between the two methods. D. Cell-wise Jaccard similarity index (JSI) of cluster marker genes reveals discrepancies with potential ramifications for cell type annotations. E-F. Element-centric consistency (ECC), represented on the colour gradient, across 100 repetitions of SLM community detection (E) and Leiden community detection (F), generated on Seurat and Monocle nearest neighbour graphs, respectively. Regions variably clustered across repetitions are revealed. G-H. JSI of CellphoneDB interaction terms, grouped by sender cluster (G) and receiver cluster (H), between Seurat clusters (rows) and Monocle clusters (columns). A one-to-one correspondence between rows and columns would indicate the same inter-cellular activity is inferred for clusters found by both methods; no such correspondence is observed in either heatmap. I-J. Pseudotime lineages inferred on the Seurat v4 (I) and Monocle v3 (J) partitions using Slingshot. The resulting trajectories depend strongly on the upstream clustering. K. The absolute difference between pseudotimes, inferred using the corresponding Seurat and Monocle pseudotimes in I and J, illustrates areas where the temporal ordering of cells varies strongly depending on the clustering results. Note the colour scale is reversed compared to other panels i.e. yellow shades indicate agreement between methods.

#### [Assessment of clustering agreement]

To quantify the similarity between the partitioning induced by the two methods, element-centric similarity (ECS) is employed (figure 4.1C); the ECS ranges from 0 to 1, with higher values indicating greater similarity in how a cell is clustered between the two methods (Supplementary Methods S2). As ECS measures clustering similarity per cell, while taking into account higher-order interactions between cells induced by the clusterings, it avoids the biases of other similarity measures with respect to cluster sizes while providing detailed information (Supplementary Methods S3). In comparing the clusterings, we note that ECS captures an intuitive notion of similarity; on the UMAP, the consistently clustered right-hand group of cells displays relatively high ECS, while the disagreed-upon central upper part has low ECS values.

For non-deterministic clustering methods, the question of stability with regard to random seed also arises i.e. assessment of the clustering variability due to the stochasticity of the clustering method. We evaluate this stability by repeatedly clustering the entire dataset with the Seurat and Monocle pipelines separately, each across 100 random seeds. Subsequently, the consistency, or frustration [48], of the clustering is evaluated by computing element-centric consistency (ECC); the ECC is computed by taking the pairwise ECS scores between all clusterings, and calculating averages per cell (Figure 4.1E-F). For the Mende et al. dataset, the Seurat pipeline overall clusters the data more consistently than the Monocle pipeline (ECC summaries: for Seurat min 0.35, median 0.96, max 1.0, stddev 0.12; for Monocle min ECC 0.33, median 0.67, max 1.0, stddev 0.17). Some regions of the UMAP, like the far right (cluster 3 in Seurat, cluster 4 in Monocle), are stable in both while others, like the lower left (clusters 4, 5, 8 in Seurat), are stable in one but not the other.

#### [Assessment of biological interpretation consequences of the assignment variability]

To further investigate the effects of clustering variability on cell type inference, we look into statistically discriminative marker genes, per cluster. These are often considered a starting point for the annotation of clusters (i.e. for assigning a biological interpretation). We compare the sets of marker genes using the Jaccard similarity index (JSI) across the ones inferred for each partitioning (figure 4.1D). Higher values for the JSI indicate greater similarity of the marker genes per cell, across both methods i.e. the cell’s identity would be interpreted similarly. We observe overall low JSI throughout the UMAP, indicating that the interpretation of the cells may vary depending on the clustering method.

Downstream from clustering and cell type identification, one common analysis is cell-cell interaction inference. Here we illustrate the outputs of CellphoneDB [38], which uses expression levels per cluster in combination with a database of ligand-receptor interactions to infer potential cellular crosstalk. This analysis is often used as a hypothesis generator, offering leads to be investigated later. CellphoneDB generates a list of interaction terms, with corresponding p-values, per pair of interacting clusters; ligands originate from a sender cluster and receptors from a receiver cluster. When applying CellphoneDB on the Mende et al. data, the inferred interaction terms differ between the Seurat and Monocle outputs. Grouping the obtained terms per sender cluster (Figure 4.1G), a binary confusion matrix, with a one-to-one relationship between rows and columns (corresponding to Seurat and Monocle clusters) would be expected if the same interactions were inferred. No such (near-perfect) relationship is observed in practice (98.6% of matrix entries 0.1 *< x <* 0.9; the 1st, 5th, 10th, 90th, 95th, 99th percentiles of the matrix entries are 0.21, 0.24, 0.27, 0.73, 0.83, 0.91, respectively). When grouping interaction terms per receiver cluster (Figure 4.1H), again no one-to-one relationship is observed (97.6% of matrix entries 0.1 *< x <* 0.9; the 1st, 5th, 10th, 90th, 95th, 99th percentiles of the matrix entries are 0.23, 0.26, 0.30, 0.80, 0.87, 0.93, respectively). This disparity in results suggests that inferred interactions, which may spur follow-up investigations, depend strongly on the upstream clustering.

For hematopoietic stem and progenitor cells, characterising differentiation trajectories is of particular interest [27]. Slingshot infers a trajectory on scRNA-seq data using cluster centroids [39]. To select start and end clusters for the inferred lineages, we used previous cell type annotations of the data (details in Methods). The resulting trajectories depend strongly on the upstream clustering (Figure 4.1I,J). In addition, using the Slingshot trajectories derived from Seurat and Monocle clusterings, a pseudotime was calculated separately on each trajectory. The resulting pseudotimes were compared per cell, revealing large differences at the top, left, and bottom of the UMAP (Figure 4.1K). This analysis illustrates that trajectory and pseudotime analyses, and resulting interpretations of differentiation dynamics, may be highly dependent on upstream clusterings.

#### [Assessment of variability on density-based clustering approaches]

A major difference between Monocle v2 vs v3 was the shift from using density-peak based clustering as default. To provide a legacy assessment, and to perform a comparison against a non graph-based clustering, we looked into the results obtained using Monocle v2 on the Mende et al. dataset. With the aim of finding the same number of clusters as Seurat, using density-peak clustering on a t-SNE dimensionality reduction yielded the partitioning of cells observed in Supplementary Figure S1B. The corresponding ECS and JSI summaries on marker genes, when comparing with the Seurat v4 clustering (Supplementary Figure S1C-D), highlight areas of similarity and difference across methods, recapitulating previous conclusions. The Slingshot trajectory, inferred from the Monocle v2 partitioning, links cell states in a biologically implausible manner (Supplementary Figure S1E). The differences between the resulting trajectories vs those based on the Seurat clustering have knock-on effects for the inferred pseudotimes (Supplementary Figure S1F). Finally, per-cluster cell-cell interaction analyses, predicted with CellphoneDB, reinforce the relatively low similarity with the Seurat outputs (Supplementary Figure S1G-H). Thus, on the Mende et al. dataset used as case study, both when comparing graph-based clustering pipelines and when comparing one graph-based and one density-peak-based pipeline, resulting hypotheses are strongly dependent on the pipeline and method in question.

### 4.2 Assessment of parameters influencing clustering results

A central task for scRNA-seq analyses is inferring cell-types; established pipelines such as Seurat [31], Monocle v3 [32], and SCANPY [17] use graph-based clustering for partitioning the data. With minor variations, the pipelines comprise 3 steps: [a] dimensionality reduction (linear or non-linear), [b] k-nearest neighbour graph inference, and [c] graph clustering. The clustering output can be significantly altered by several parameters (illustrated in 4.2A). Since algorithms across all steps are stochastic, we first evaluated the effect of using different random seeds, by calculating the EC consistency over 100 runs (i.e. the resulting partitions). As shown in 4.2B the random seeds do have an effect on the stability of the output (ECC summary: min 0.38, mean 0.90, median 0.97, max 1.0, stdev 0.12); the corresponding resulting partitions and their frequencies are overviewed in Supplementary Figure S2.1; for this dataset, using default parameters there is no high-frequency partition, however we also note that some of the changes in cluster assignation are minimal. Across all subsequent analyses the stability of results is assessed, and summarised over 100 random seeds.

#### [Dimensionality reduction]

Focusing first on the dimensionality reduction step, we analyzed the inputs, the parameters and algorithmic details that influence the PCA outputs. The input set (feature set) is crucial for calculating the distribution of cells in the reduced space. Four categories of feature sets frequently used in practice are the all genes or most abundant genes (default in Monocle [49, 41]), highly variable genes (default in Seurat [50, 16]), and the intersection between the most abundant and the highly variable genes [41]. In 4.2C we summarise the ECC distributions and note the differences in variability; the set of most abundant features tends to drive tighter distributions, with an ECC converging to 1 (i.e. no variability) as the size of the set increases (ECC summaries: for PCA with all genes min 0.96, mean 0.99, median 1.00, max 1.0, stdev 0.001; for UMAP with all genes min 0.37, mean 0.90, median 0.94, max 1.0, stdev 0.10). More variability is observed for the highly variable set of features (ECC summaries: for PCA with 3000 genes min 0.51, mean 0.92, median 0.96, max 1.0, stdev 0.09; for UMAP with 3000 genes min 0.34, mean 0.80, median 0.84, max 1.0, stdev 0.15); a middle ground, in terms of ECC variability is observed for the intersection of most abundant and most variable features (ECC summaries: for PCA with 300 genes min 0.50, mean 0.93, median 0.99, max 1.0, stdev 0.11; for UMAP with 300 genes min 0.26, mean 0.85, median 0.88, max 1.0, stdev 0.15). In addition, the underlying embedding influences the stability; the UMAP embedding (represented in blue in 4.2C) leads to lower and more variable ECC than the PCA one. The purpose of the incremental approach was to determine the largest subset of genes not affected by noise. For the PCA embedding we note an increase of variability for 2000+ most abundant features; for the UMAP embedding little variability is observed for the <2000 HV features, followed by a reduction for 2000+ genes (Supplementary Figures S2.2, S2.3).

Yet another technical parameter, that influences the stability of the PCA results, is the tolerance. In practice, approximate PCA calculations are often used for runtime efficiency at a minimal cost on precision; the Monocle and Seurat pipelines rely on the `irlba` package [43] with default values for the tolerance set at 1e-5. The user-defined threshold for the tolerance impacts the stability of the results, through error propagation; in 4.2D we illustrate the PC differences obtained for tolerance thresholds of 1e-5, 1e-10 and 1e-15 especially noticeable on the last PCs (differences are calculated against the `prcomp` calculation, precise up to machine precision); additional PCs are presented in Supplementary Figure S2.4.

#### [Graph construction]

The building of the graph relies on the kNN algorithm [51]; therefore the number of neighbours will have an impact on the partitioning. In panels 4.2F, G we illustrate the co-variation between the number of neighbours (x-axis) and the number of clusters (k) and the EC consistency, respectively. Panels 4.2E and F underline that an increase in the number of neighbours leads to a better connected graph; the number of connected components acts as a lower bound for the number of clusters. Individual clusters can still be sub-partitioned into finer populations. Panel 4.2F also underlines the effect of two other parameters: the embedding and the graph type (unweighted vs weighted). The former contrasts the PCA vs UMAP embedding, with the UMAP systematically producing more connected components (for the Cuomo et al. dataset twice as many) than the PCA. The latter focuses on the edge weights of the resulting graph. The shared nearest neighbor (SNN) weighted version, used in Seurat and Monocle, results in more stable partitions across different seeds (Figure 4.2G); moreover a better stability is obtained in comparison with the NN version as well. The differences in the SNN implementation between the Seurat and the Monocle pipelines are detailed in Supplementary Information. This summary panel also recapitulates the differences between UMAP and PCA-based graphs; for fewer neighbours, the ECC difference is significant, however the increase in the number of neighbours evens out the comparison (ECC summary when using 30 number of neighbours: for SNN based on PCA, min 0.82, mean 0.99, median 0.99, max 1.0, stdev 0.008; for NN based on PCA, min 0.46, mean 0.91, median 0.96, max 0.99, stdev 0.11; for SNN based on UMAP, min 0.26, mean 0.85, median 0.88, max 1.0, stdev 0.15; for NN based on UMAP, min 0.26, mean 0.84, median 0.86, max 1.0, stdev 0.13); an overview of areas of lower consistency is presented in Supplementary Figure S2.5.

**Figure 4.2:**
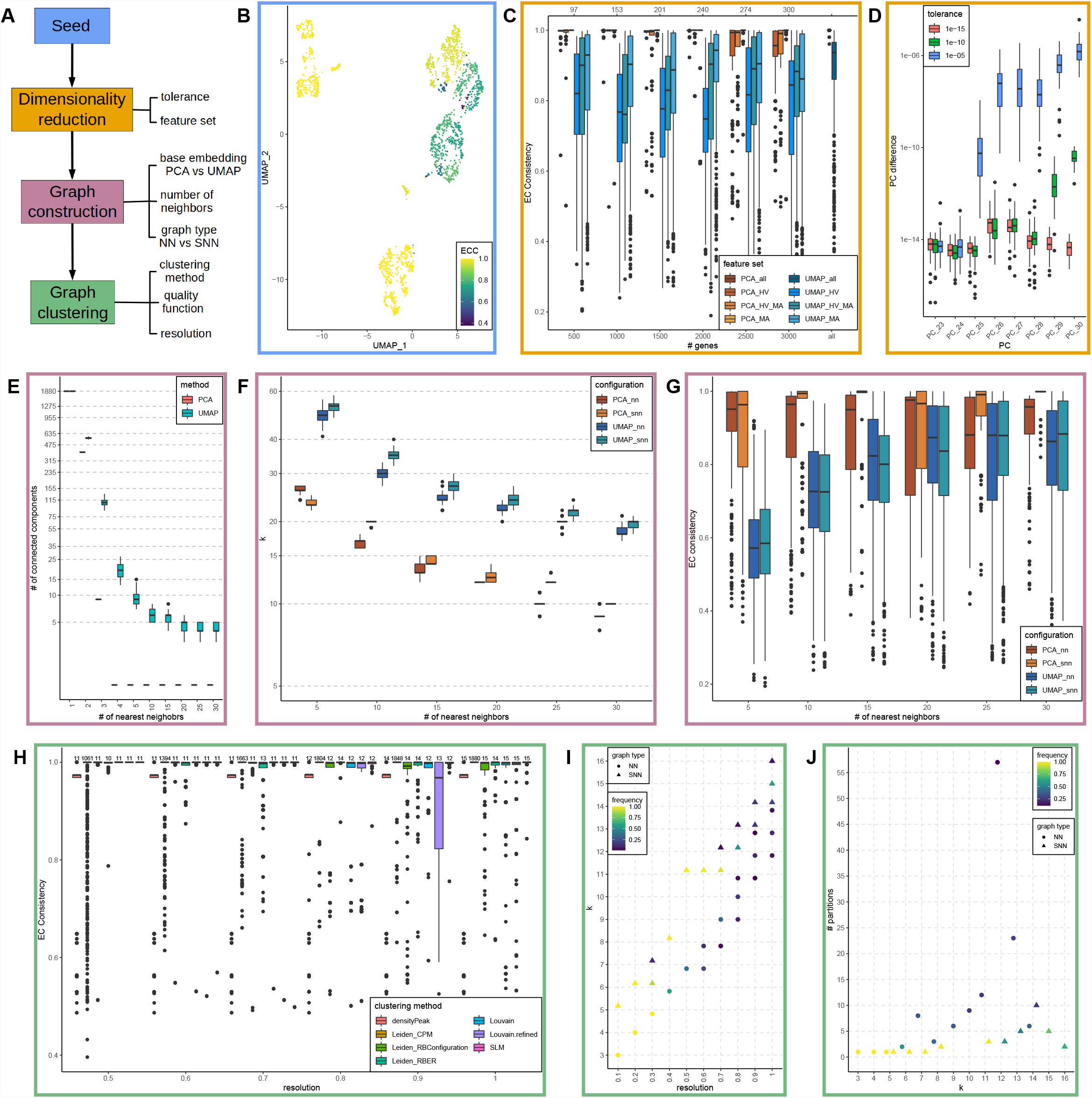
Stability of graph-based clustering pipeline parameters across random seeds. The analysis is performed on the Cuomo et al. SMART-seq dataset. A. Diagram underlining the steps of the clustering pipeline (common to e.g. Seurat and Monocle) and the corresponding main parameters that drive the variability. B. UMAP representation of ECC, summarising the variability across 100 different random seeds. C. Summary of the ECC distributions (over 100 runs) for various feature sets (most abundant genes, highly variable genes and their intersection), various numbers of genes considered (500 to 3000, in increments of 500) and on a PCA or UMAP base embedding used for building the graph (orange and blue gradients, respectively). D. Summary of the distributions of differences (over 100 runs) across components for the precise vs the approximate (irlba) calculation of the PCA. The colour scheme corresponds to typical precision levels. E. Summary of the co-variation (over 100 runs) between the number of nearest neighbours and number of connected components; the differences induced by the embedding (PCA and UMAP) are underlined by colour. F. Summary of the co-variation (over 100 runs) of the number of the nearest neighbours vs number of clusters; the differences induced by the base embedding and the graph type (NN: unweighted; SNN: weighted) are underlined by colour. G. Summary of ECC distributions (over 100 runs) induced by the graph type (NN vs SNN); the differences induced by the base embedding are underlined by colour. H. Summary of ECC distributions (over 100 runs) induced by the clustering method vs the resolution parameter (x-axis). Above each boxplot the number of clusters for the most frequent partition is presented. I. Summary of co-variation between the resolution parameter and the number of clusters. The point shape indicates the graph type, the colour gradient is proportional to the frequency of the most common partition. J. Summary of the stability of partitioning (over 100 runs), on fixed number of clusters, evaluated on the number of resulting partitions (y-axis); the colour gradient is proportional to the frequency of the most common partition.

#### [Graph clustering]

Community detection or graph clustering methods were adopted by the community due to their increased performance and ability to adapt to size and shape characteristics of clusters. These were proposed in clustering pipelines such as PhenoGraph [52] and became default choices for single-cell pipelines (Seurat, Monocle). With a bottom-up approach, the methods aim to form clusters that optimise an objective function, with convergence reached after several iterations. The longstanding Louvain algorithm [45] was superseeded by improved methods such as Louvain with multi-level refinement [46] or SLM [34]; the current improvement is the Leiden algorithm [35].

To assess the stability of results with respect to the clustering method, we illustrate in Figure 4.2H the ECC distributions corresponding to the four approaches (all available in the Seurat package; only Louvain and Leiden are available in Monocle). Also contrasted is densityPeak, a density-based clustering method used as default in Monocle v2. (in Supplementary Figure S2.6 we also included the k-means with random initialisation, for theoretical consistency). The stability does not significantly differ across methods, although Louvain refined performs noticeably worse for resolution=0.9 (ECC summary: for SLM, min 0.51, mean 0.99, median 0.99, max 1.0, stdev 0.02; for densityPeak min 0.49, mean 0.96, median 0.97, max 1.0, stdev 0.07; for Leiden RBConfiguration min 0.55, mean 0.97, median 0.99, max 1.0, stdev 0.07; for Louvain min 0.59, mean 0.97, median 0.99, max 1.0, stdev 0.04; for Louvain refined min 0.42, mean 0.92, median 0.97, max 1.0, stdev 0.11). Also assessed is the difference between objective functions (quality functions or metrics) optimised by a graph-based clustering algorithm, namely Constant Potts model (CPM), Reichardt and Bornholdt’s Potts model with a configuration null model (RBConfiguration), and Reichardt and Bornholdt’s Potts model with an Erdős-Rényi null model (RBER) [53] (Figure 4.2H). For the same resolution, the quality functions may lead to a different number of clusters (e.g., for resolution of 0.7, CPM leads to 1663 clusters, RBConfiguration 11 and RBER 13). The CPM quality function used as default in Monocle, bounds the cluster size based on the clustering resolution and number of nearest neighbours (Supplementary Figure S3). These bounds can partly explain the increased number of clusters obtained when using CPM rather than RBConfiguration and RBER, while keeping the other parameters constant.

Lastly, an essential parameter that directly controls the number of communities is the resolution. The co-variation between the resolution and the number of clusters is summarised in Figure 4.2I (higher values of the resolution lead to an increase in the number of clusters). This summary also provides an overview of suitable (stable) ranges for the resolution parameter and reviews the overall stability of the partition on random seeds; the colour gradient is proportional to the number of occurrences of the most frequent partition. Another detail embedded in the panel is the effect/ difference between the two graph types; the number of clusters is smaller for the NN graph compared to the SNN. In addition, the number of stable resolution - number of cluster correspondences is higher for the SNN (7 entries have a frequency >0.85) compared with NN (3 entries have a frequency >0.85).

Another important characteristic is the number of partitions (the co-variation between the stability of the number of clusters and the number of resulting, different partitions is summarised in Figure 4.2J). A high number of different partitions implies a lower stability for a given number of clusters. The colour gradient is proportional to the frequency of the most common partition for a fixed number of clusters. A high number of different partitions may be acceptable if the frequency of the most common one is close to 1. Conversely, a high number of partitions, each with low frequency, indicates high instability. The graph-type detail is also embedded in the panel; similarly as in panel I, NN graphs are more unstable compared to SNN graphs (the maximum number of different partitions for NN is 57, compared with 10 for SNN). The conclusions also hold for a wider range of resolutions, and with different ECS thresholds (Supplementary Figures S2.7, S2.8, S2.9).

### 4.3 Clustering results vary across data modalities

As new assays become available for high-throughput profiling of multiple modalities at single-cell resolution, new analysis-driven challenges emerge [54]. As with individual single-cell modalities (e.g. scRNA-seq) the grouping of cells into clusters is central to multimodal analyses [55]. Recently developed approaches for multimodal clustering draw on the information available in each modality to calculate the partitioning of cells [14, 15]. These approaches rely on the assumption that there exists an underlying cell grouping that could be retrieved regardless of modality, and using information across modalities will help recover this latent partitioning. The agreement across individual modalities was considered by Gao et al. [56], who developed a statistical test of independence between clusterings. The test was initially applied on the Pioneer 100 Wellness Project dataset of multiple data types, across several timepoints, on a cohort of participants; the authors concluded that clusterings derived from different modalities were not dependent for several pairwise comparisons. While their statistical test fits Gaussian mixture models to each data modality to test for independence, ECS compares concrete partitions. In addition, ECS offers a per-observation measure of the extent to which modality-specific clusterings agree, providing a more detailed picture of the similarities across modalities (Supplementary Methods S2).

To assess the applicability of ClustAssess on multiomics data, we present a case study on a single-cell RNA and ATAC dataset of peripheral blood provided by 10x Genomics. First, we consider each modality separately; to obtain representative results, we construct nearest neighbour graphs separately for each modality and partition 100 times across random seeds with SLM clustering. We select the most common partition, for each modality, for downstream analyses. We identify 15 clusters for the RNA modality and 11 for the ATAC one (Figure 4.3A,B). Using ECS we underline the differences between the partitions generated on each modality (Figure 4.3D; ECS summary: min 0.0, max 0.96, median 0.47, mean 0.47). The two isolated islands in the upper part of the UMAP display good agreement, as they each make up one cluster in both clusterings. Conversely, the two larger bodies of cells towards the bottom of the UMAP display lower ECS scores overall, underlining the variability in how the cells are

**Figure 4.3:**
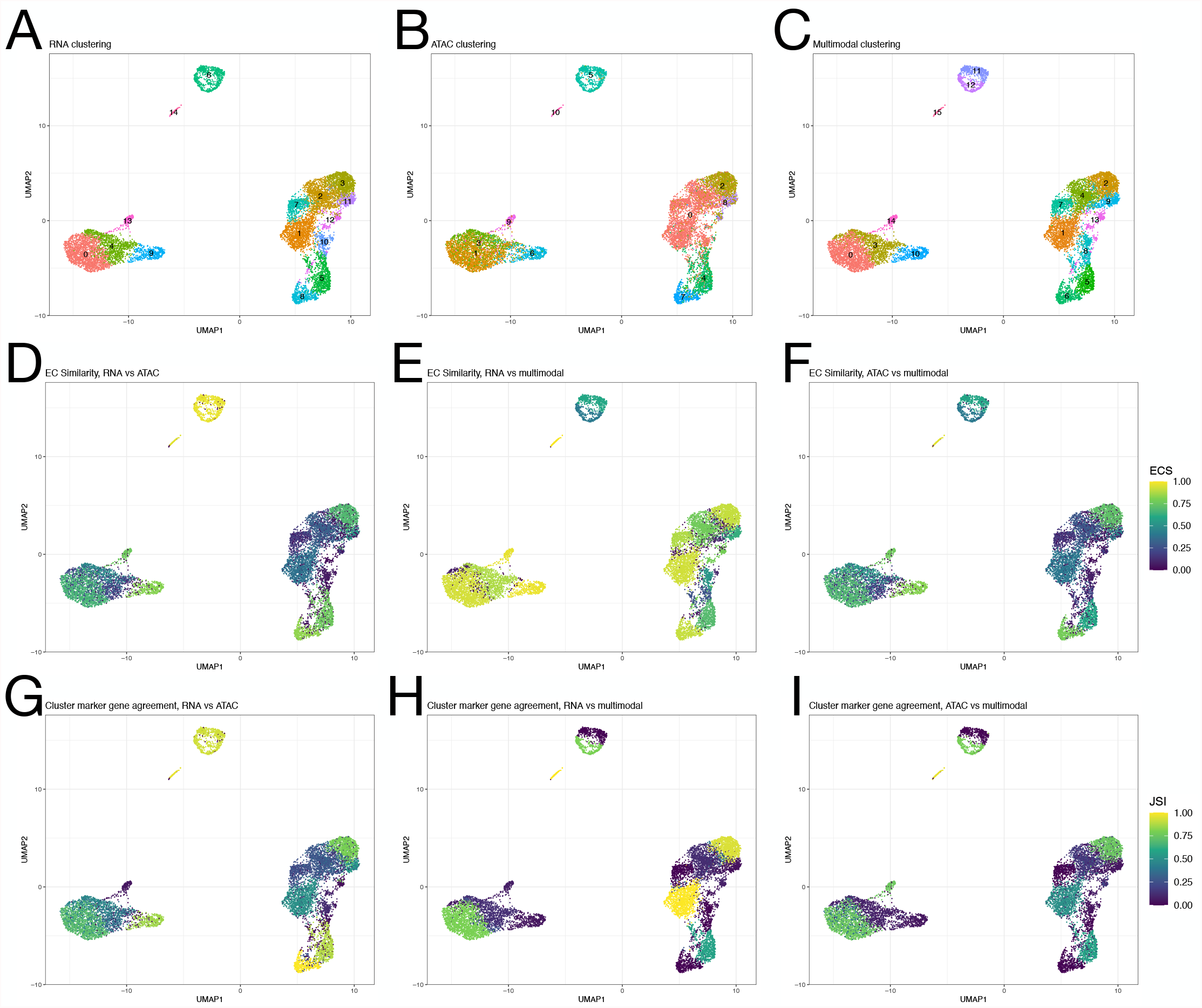
Differences between clustering results created on different modalities of single-cell RNA+ATAC data. A-C. Partitions obtained using SLM community detection on nearest neighbour graphs generated on PCA reduction of RNA modality (A), LSI reduction of ATAC modality (B), and on the weighted nearest neighbour graph created jointly on the PCA and LSI reductions (C). Each graph was clustered 100 times across random seeds; the most frequent partition for each modality is presented (with frequency 0.12, 0.13, 0.09 for RNA, ATAC, and joint RNA+ATAC partitions, respectively). D-F. Element-centric similarity (colour gradient) comparing RNA and ATAC clusterings (D), RNA and joint clustering (E), ATAC and joint clustering (F). Differences are observed between the RNA and ATAC clusterings, while the RNA and joint clusterings display higher agreement. G-H. Jaccard similarity index (JSI) of cluster marker genes obtained using the partitionings on each modality. Higher JSI indicates better agreement between markers across modalities, which may have implications for downstream annotations of cellular identity. All results visualised on UMAP generated on PCA from RNA modality of the data. partitioned depending on modality.

When considering a multimodal clustering using information from both RNA and ATAC modalities, we obtain 16 clusters (Figure 4.3C). Notably, the uppermost island of cells, grouped into a single cluster in both RNA and ATAC modalities, is now split into two parts. Using ECS to evaluate the per-cell agreement between the multimodal clustering with each of the unimodal clusterings (Figure 4.3E,F), we observe overall high similarity between the multimodal and the RNA clusterings (ECS summary: min 0.0, max 0.99, median 0.87, mean 0.76), while the agreement between multimodal and ATAC modalities is lower (ECS summary: min 0.0, max 0.95, median 0.41, mean 0.44), and broadly similar to the agreement between RNA and ATAC, apart from the top island (4.3D). Confusion matrices of the partitions for each modality underline the discrepancies between the clusterings (Supplementary Figure S4A-C). We also assessed the clustering stability on random seeds, on each modality. Across 100 random seeds, ECC distributions reveal high stability overall, and suggest the ATAC modality is most consistently clustered, while the RNA and joint RNA+ATAC modalities have some areas of inconsistently clustered cells; these areas are generally disjoint on the UMAP across the modalities (Supplementary Figure S4D-F).

To observe how the differences in clustering across modalities may affect downstream interpretation, we calculate the most discriminative marker genes, per cluster, for each partitioning. Comparing the marker genes across modalities using the JSI approach, we identify a mixture of areas of both higher and lower agreement (Figure 4.3G-I). Notably, the top island has high JSI for RNA vs ATAC, but lower when comparing against the multimodal clustering. Overall, the patterns of JSI recapitulate the ECS results obtained when comparing each unimodal clustering against the multimodal one.

Given the conceptual differences between the measured assays and the discrepancies between the RNA and ATAC clusterings, it is worth considering whether there is indeed an underlying grouping of cells that will be reflected in every modality. If no such universal grouping is visible and each modality reflects a distinct pattern of cell groups, attempts at pooling information across modalities during clustering, as we have done here, may be unwise. For this specific dataset, the multimodal clustering draws heavily on the RNA component, while the contribution from the ATAC modality is less significant; ECS quantifies this co-variation for each region of the UMAP.

## 5 Discussion

### 5.1 ClustAssess identifies robust, dataset-specific parameter ranges

The outputs of each step of the clustering pipeline are influenced by user-defined parameters. The main aim of `ClustAssess` is to evaluate the robustness of inferred partitions to technical variations (e.g. changes in random seed) to ensure a reflection of true biological signal. The assessment of the stability is built on the ECC across partitions generated with different random seeds. A side effect of this analysis is also the identification of “optimal” ranges, and the characterisation of complex co-variations, for the various parameters used throughout the clustering pipeline. This alternative use of `ClustAssess` is exemplified in an additional vignette provided as part of the package.

We first focus on the dimensionality reduction step; an essential parameter is the feature set. The user can provide a feature set, including frequently used options such as the most abundant genes [49, 41], or highly variable genes [50, 16], or other custom subset specified by the user [57]. Also assessed are different sizes (number of entries) of the feature set; this incremental approach is essential for determining the largest subset of genes unaffected by noise [22, 21] which can alter downstream interpretations. The stability of each feature set is summarised using boxplots; the distribution of the ECC on the reduced UMAP space provides additional insights on the localisation of unstable regions on the UMAP topography.

Next, several parameters involved in the graph construction phase are evaluated: [a] the base embedding i.e. the dimensionality reduction of the input data on which the graph is built, [b] the number of neighbours and [c] the graph type i.e. unweighted on nearest neighbours, or weighted using shared nearest neighbours. In turn, these parameters are assessed with respect to the graph connectivity (number of connected components). While PCA embeddings are widely used, nonlinear dimensionality reductions including UMAP are popular in single-cell data analysis for their ability to preserve local structure without overcrowding the visualisation [58]. Increasing the number of neighbours leads to a decrease in the number of connected components (more neighbours result in a better connected graph); the number of connected components provides a lower bound for the number of clusters obtainable by downstream community detection methods. The concept of shared nearest neighbours was adapted to the single cell setting in [52]. Changing the approach from nearest neighbours to shared nearest neighbours leads to differences such as: the SNN approach adds weights to the graph using the JSI between neighbourhoods of cells and increases the connectivity by introducing edges between cells that are indirect neighbours. Also assessed is the relationship between graph construction parameters and the number of communities that a graph clustering algorithm will delineate.

The central component of the clustering pipeline is the clustering method; we evaluate the impact of frequently used community detection algorithms in the context of varying/ optimising the resolution parameter. We focus on four methods available in Seurat: Louvain [45], Louvain with multilevel refinement [46], SLM [34] and Leiden [35]. The resolution parameter is critical for determining the final number of clusters (increasing the resolution results in a higher number of partitions) and in the subsequent biological interpretation. The stability of results is assessed across a gradient of resolutions versus the number of clusters in summary plots (as presented in Figure 4.2I) showcasing the frequency of the most stable partition and the number of resulting partitions as well as using the ECCs on UMAP representations.

### 5.2 Aligning the Seurat and Monocle pipelines by adjusting and assessing step-wise parameters

The two most frequently used R-based single-cell pipelines, Seurat [31] and Monocle [32], share similar steps for processing the data i.e. normalisation of expression levels, linear and non-linear dimensionality reduction, graph building based on *k* nearest neighbours, and community detection on the resulting graph. However, as observed in the 10x and SMART-seq case studies presented in the Results section, there are two main sources of variability: [a] the default (and occasionally optimised) parameter values/ choices lead to different results, and potentially different biological interpretations and [b] the stochastic components of the methods may introduce significant variability e.g. in UMAP topologies (Supplementary Figure S5 panels S1-2, M1-2). Both sources of variability are technical; to address the reproducibility of results, we focused on the adjustment of parameters to ensure identical outputs. Each pipeline uses a specific set of default values; thus, to obtain identical outputs, it is necessary to match the values. The comparison is performed step-wise (Supplementary Figure S5).

The first task is achieving identical representations on a dimensionality reduction; Seurat and Monocle rely on different sets of features for the PCA dimensionality reduction i.e. Monocle uses all genes [32], while Seurat focuses only the highly variable ones [31]. The underlying feature set influences the topography (Supplementary Figure S5 panels S3,4 and M3,4); the all genes set induces a partitioning into numerous small islands (Supplementary Figure S5 panels S4, M3), whereas the highly variable set of genes produces more compact groups (Supplementary Figure S5 panels S3, M4). Moreover, even though UMAP is the default non-linear representation in both pipelines, default values for the min_dist and n_neighbors parameters differ (S5 S5-8, M5-8), and impact the embedding. Using identical values for `min_dist` and `n_neighbors` (0.3 and 30 respectively) results in identical dimensionality reductions.

Next, the graph is built on a reduced space embedding; by default, Seurat uses the Principal Components as a base embedding [31], while Monocle uses UMAP [32]. Different bases induce a variable number of clusters (Supplementary Figure S5 panels S9 - S10 and M9 - M10). Yet another difference stems from the weighting of graphs (Monocle uses as default NN, Seurat uses as default SNN). The graph type influences the number of clusters and the cells assigned to them (Supplementary Figure S5 panels S11 and S12); to achieve agreement we set these two parameters to PCA and SNN respectively. It is worth noting that Seurat and Monocle have a slightly different implementation of the graph construction method, thus achieving identical graphs requires an additional processing of the results (further details in Supplementary Information).

The pipeline concludes with applying a graph clustering algorithm (Supplementary Figure S5 panels S13-18 and M13-18). To assess the comparability of results, we applied the same clustering methods, yet some differences were observed (Supplementary Figure S5 panels S13, S14, M13, M14). The community detection algorithms delineate clusters by optimising an objective function [35], and changing objective functions leads to a different final partitioning (Supplementary Figure S5 panels S15, S16, M15, M16). The range of values for the resolution parameter impacting the clustering output is highly dependant on the objective function (i.e. the resolution is incorporated differently in different objective functions). In addition, the resolution parameter directly impacts the final number of clusters (Supplementary Figure S5 panels S17, S18, M17, M18); for the CPM quality function in particular, the combination of resolution and number of neighbours provides tight bounds on the size of communities [53], which may be overly restrictive for the context of subpopulation identification in single-cell data (Supplementary Figure S3). Given the iterative nature of the graph clustering methods, the convergence of the graph clusters depends on the number of iterations. Seurat and Monocle use as default different number of iterations (10 and 2, respectively), which generate slightly different results (Supplementary Figure S5 panel SM19). Using the same value (10) leads to identical partitions(Supplementary Figure S5 panels S20, M20 and SM21).

We thus illustrated that Monocle v3 and Seurat v4 do lead to identical partitions if the corresponding parameters are set to the same values. We affirmed that the observed differences are purely down to choices of parameters and underlined the importance of parameter optimisation to ensure robust and reproducible partitions (and downstream biological interpretations).

These results are recapitulated on earlier versions of Seurat; the only difference in the clustering pipeline between v3 and v4 of Seurat focuses on the `FindNeighbors` method (used for graph construction). The default package used for computing the nearest neighbours changed from rann to annoy, but Seurat v4 accommodates both options. Our previous conclusions hold for the Monocle v3 vs Seurat v3 comparison.

### 5.3 Scalability of the data-driven assessment of parameters

To assess the scalability of ClustAssess components, with a particular focus on the stability assessment pipeline, we performed time benchmarks across variable numbers of cells; the results were obtained by applying the pipeline on variable-size subsamples of cells of the Mende et al. dataset; the subsamples were generated using geometric sketching [59]. The number of iterations for parameter assessment functions was set to 30 (the function calls were similar to the ones presented in the vignette).

The `clustering_importance, feature_stability` and `nn_importance` scale similarly and monotonically as the number of cells increases (Supplementary Figure S6A,B). The `resolution_importance` steps performs better, although performance is still superlinear. `nn_n_conn_comps` is the only method scaling sublinearly with the number of cells, which may be explained by the absence of calls to clustering functions. Little variability is observed for different values of the ECS threshold (Supplementary Figure S6A,B).

The performance gain associated with using multiple cores increases with the size of the dataset (Supplementary Figure S6B)); however for small datasets the additional overhead may lead to an increase in runtime (e.g. `resolution_importance` and `clustering_importance` achieve minimal runtime, on 1319 cells, on only one core, Supplementary Figure S6B). Using smaller subsets of cells on the benchmark data, the sequential run was faster than the parallelised R instances, which need to copy the variables and collate the results obtained on individual threads. Another context when multiple cores increase the runtime is observed for a small number of iterations.

The performance of all methods is highly dependent on the characteristics of the dataset (e.g. number of cells, sequencing depth), the number of repetitions and the values of the parameters selected throughout the clustering pipeline (e.g. UMAP calculations are slower than PCA, and Louvain is a faster clustering method than SLM). The package documentation and vignettes provide further guidance on usage and parameter choices, and additionally illustrate how to integrate ClustAssess with the Seurat single-cell toolkit [31].

## 6 Conclusion

Despite their shortcomings, ARI and other measures of clustering similarity are widely used in benchmarks, either for assessing clustering methods themselves [25, 60], or to gauge the consequences of upstream decisions such as batch correction and feature selection [41, 61]. In these studies, partitions are compared both against ground-truth clusterings and across methods, parameters, etc. Through `ClustAssess`, we make ECS readily available for the R community, as a more reliable and flexible alternative when evaluating clustering similarity. Moreover, progressing from a pairwise view (PAC, ARI) to an element-wise view (ECS) on stability and similarity enables the understanding of clustering robustness at a finer level of resolution than before. In this work, its efficiency was exemplified from different angles that assist with distinguishing between technical variability and robust and reproducible biological signal. We illustrated the impact of (non-biological) technical factors, from parameter choices to random seeds, on single-cell partitionings, and their downstream effects on the biological conclusions. Through `ClustAssess`, we enable quantitative evaluations of technical effects on single-cell analyses, and assist users with the identification of parameter ranges that lead to more robust analyses.

## Author contributions

AS and IM designed the study, IM, AS supervised the study. All authors contributed to the implementation of the R package, to all analyses, and wrote and revised the manuscript. All authors read and approved the submitted manuscript.

## Acknowledgements

This research was funded by the Wellcome Trust [203151/Z/16/Z] and the UKRI Medical Research Council [MC_PC_17230]. For the purpose of open access, the authors applied a CC BY public copyright licence to all versions of the manuscript arising from this submission.

We thank the Core Bioinformatics team at the Cambridge Stem Cell Institute for constructive discussions and feedback during the preparation of the manuscript. AM thanks Prof Liviu Ciortuz, Faculty of Computer Science, Alexandru Ioan Cuza University, Iasi, Romania, for supervision and feedback.

## References

[1] Valentine Svensson, Eduardo da Veiga Beltrame, and Lior Pachter. A curated database reveals trends in single-cell transcriptomics. Database, 2020(baaa073), January 2020.

[2] Daniel Dominguez, Yi-Hsuan Tsai, Nicholas Gomez, Deepak Kumar Jha, Ian Davis, and Zefeng Wang. A high-resolution transcriptome map of cell cycle reveals novel connections between periodic genes and cancer. Cell Research, 26(8):946–962, August 2016.

[3] Jeffrey A. Farrell, Yiqun Wang, Samantha J. Riesenfeld, Karthik Shekhar, Aviv Regev, and Alexander F. Schier. Single-cell reconstruction of developmental trajectories during zebrafish embryogenesis. Science, 360(6392), June 2018.

[4] Zinaida Good, Jolanda Sarno, Astraea Jager, Nikolay Samusik, Nima Aghaeepour, Erin F. Simonds, Leah White, Norman J. Lacayo, Wendy J. Fantl, Grazia Fazio, Giuseppe Gaipa, Andrea Biondi, Robert Tibshirani, Sean C. Bendall, Garry P. Nolan, and Kara L. Davis. Single-cell developmental classification of B cell precursor acute lymphoblastic leukemia at diagnosis reveals predictors of relapse. Nature Medicine, 24(4):474–483, April 2018.

[5] Runxia Liu, Yang-Hui Jimmy Yeh, Ales Varabyou, Jack A. Collora, Scott Sherrill-Mix, C. Conover Talbot, Sameet Mehta, Kristen Albrecht, Haiping Hao, Hao Zhang, Ross A. Pollack, Subul A. Beg, Rachela M. Calvi, Jianfei Hu, Christine M. Durand, Richard F. Ambinder, Rebecca Hoh, Steven G. Deeks, Jennifer Chiarella, Serena Spudich, Daniel C. Douek, Frederic D. Bushman, Mihaela Pertea, and Ya-Chi Ho. Single-cell transcriptional landscapes reveal HIV-1–driven aberrant host gene transcription as a potential therapeutic target. Science Translational Medicine, 12(543), May 2020.

[6] David Lähnemann, Johannes Köster, Ewa Szczurek, Davis J. McCarthy, Stephanie C. Hicks, Mark D. Robinson, Catalina A. Vallejos, Kieran R. Campbell, Niko Beerenwinkel, Ahmed Mahfouz, Luca Pinello, Pavel Skums, Alexandros Stamatakis, Camille Stephan-Otto Attolini, Samuel Aparicio, Jasmijn Baaijens, Marleen Balvert, Buys de Barbanson, Antonio Cappuccio, Giacomo Corleone, Bas E. Dutilh, Maria Florescu, Victor Guryev, Rens Holmer, Katharina Jahn, Thamar Jessurun Lobo, Emma M. Keizer, Indu Khatri, Szymon M. Kielbasa, Jan O. Korbel, Alexey M. Kozlov, Tzu-Hao Kuo, Boudewijn P.F. Lelieveldt, Ion I. Mandoiu, John C. Marioni, Tobias Marschall, Felix Mölder, Amir Niknejad, Lukasz Raczkowski, Marcel Reinders, Jeroen de Ridder, Antoine-Emmanuel Saliba, Antonios Somarakis, Oliver Stegle, Fabian J. Theis, Huan Yang, Alex Zelikovsky, Alice C. McHardy, Benjamin J. Raphael, Sohrab P. Shah, and Alexander Schönhuth. Eleven grand challenges in single-cell data science. Genome Biology, 21(1):31, February 2020.

[7] Ricard Argelaguet, Anna S. E. Cuomo, Oliver Stegle, and John C. Marioni. Computational principles and challenges in single-cell data integration. Nature Biotechnology, pages 1–14, May 2021.

[8] Vladimir Yu Kiselev, Tallulah S. Andrews, and Martin Hemberg. Challenges in unsupervised clustering of single-cell RNA-seq data. Nature Reviews Genetics, 20(5):273–282, May 2019.

[9] Tallulah S. Andrews and Martin Hemberg. Identifying cell populations with scRNASeq. Molecular Aspects of Medicine, 59:114–122, February 2018.

[10] Wouter Saelens, Robrecht Cannoodt, Helena Todorov, and Yvan Saeys. A comparison of single-cell trajectory inference methods. Nature Biotechnology, 37(5):547–554, May 2019.

[11] Rongxin Fang, Sebastian Preissl, Yang Li, Xiaomeng Hou, Jacinta Lucero, Xinxin Wang, Amir Motamedi, Andrew K. Shiau, Xinzhu Zhou, Fangming Xie, Eran A. Mukamel, Kai Zhang, Yanxiao Zhang, M. Margarita Behrens, Joseph R. Ecker, and Bing Ren. SnapATAC: A Comprehensive Analysis Package for Single Cell ATAC-seq. bioRxiv, page 615179, August 2020.

[12] Lukas M. Weber and Mark D. Robinson. Comparison of clustering methods for highdimensional single-cell flow and mass cytometry data. Cytometry Part A, 89(12):1084–1096, 2016.

[13] Laura Capolupo, Irina Khven, Luigi Mazzeo, Galina Glousker, Francesco Russo, Jonathan Paz Montoya, Sylvia Ho, Dhaka R. Bhandari, Andrew P. Bowman, Shane R. Ellis, Romain Guiet, Johannes Muthing, Bernhard Spengler, Ron M. A. Heeren, Gian Paolo Dotto, Gioele La Manno, and Giovanni D’Angelo. Sphingolipid Control of Fibroblast Heterogeneity Revealed by Single-Cell Lipidomics. bioRxiv, page 2021.02.23.432420, February 2021.

[14] Ricard Argelaguet, Britta Velten, Damien Arnol, Sascha Dietrich, Thorsten Zenz, John C Marioni, Florian Buettner, Wolfgang Huber, and Oliver Stegle. Multi-Omics Factor Analysis—a framework for unsupervised integration of multi-omics data sets. Molecular Systems Biology, 14(6):e8124, June 2018.

[15] Joshua D. Welch, Velina Kozareva, Ashley Ferreira, Charles Vanderburg, Carly Martin, and Evan Z. Macosko. Single-Cell Multi-omic Integration Compares and Contrasts Features of Brain Cell Identity. Cell, 177(7):1873–1887.e17, June 2019.

[16] Tim Stuart, Andrew Butler, Paul Hoffman, Christoph Hafemeister, Efthymia Papalexi, William M. Mauck, Yuhan Hao, Marlon Stoeckius, Peter Smibert, and Rahul Satija. Comprehensive Integration of Single-Cell Data. Cell, 177(7):1888–1902.e21, June 2019.

[17] F. Alexander Wolf, Philipp Angerer, and Fabian J. Theis. SCANPY: Large-scale single-cell gene expression data analysis. Genome Biology, 19(1):15, February 2018.

[18] Cole Trapnell, Davide Cacchiarelli, Jonna Grimsby, Prapti Pokharel, Shuqiang Li, Michael Morse, Niall J. Lennon, Kenneth J. Livak, Tarjei S. Mikkelsen, and John L. Rinn. The dynamics and regulators of cell fate decisions are revealed by pseudotemporal ordering of single cells. Nature Biotechnology, 32(4):381–386, April 2014.

[19] Tamim Abdelaal, Lieke Michielsen, Davy Cats, Dylan Hoogduin, Hailiang Mei, Marcel J. T. Reinders, and Ahmed Mahfouz. A comparison of automatic cell identification methods for single-cell RNA sequencing data. Genome Biology, 20(1):194, September 2019.

[20] Stephanie C Hicks, F William Townes, Mingxiang Teng, and Rafael A Irizarry. Missing data and technical variability in single-cell RNA-sequencing experiments. Biostatistics, 19(4):562–578, October 2018.

[21] Gökcen Eraslan, Lukas M. Simon, Maria Mircea, Nikola S. Mueller, and Fabian J. Theis. Single-cell RNA-seq denoising using a deep count autoencoder. Nature Communications, 10(1):390, January 2019.

[22] Ilias Moutsopoulos, Lukas Maischak, Elze Lauzikaite, Sergio A Vasquez Urbina, Eleanor C Williams, Hajk-Georg Drost, and Irina I Mohorianu. noisyR: Enhancing biological signal in sequencing datasets by characterizing random technical noise. Nucleic Acids Research, 49(14):e83, August 2021.

[23] Eleanor C. Williams, Ruben Chazarra-Gil, Arash Shahsavari, and Irina Mohorianu. The sum of two halves may be different from the whole. Effects of splitting sequencing samples across lanes, November 2021.

[24] Aaron T.L. Lun, Davis J. McCarthy, and John C. Marioni. A step-by-step workflow for low-level analysis of single-cell RNA-seq data with Bioconductor. F1000Research, 5:2122, October 2016.

[25] Angelo Duò, Mark D. Robinson, and Charlotte Soneson. A systematic performance evaluation of clustering methods for single-cell RNA-seq data. F1000Research, 7:1141, November 2020.

[26] Saskia Freytag, Luyi Tian, Ingrid Lönnstedt, Milica Ng, and Melanie Bahlo. Comparison of clustering tools in R for medium-sized 10x Genomics single-cell RNA-sequencing data. F1000Research, 7:1297, December 2018.

[27] Nicole Mende, Hugo P Bastos, Antonella Santoro, Krishnaa T Mahbubani, Valerio Ciaurro, Emily Francesca Calderbank, Mariana Quiroga Londoño, Kendig Sham, Giovanna Mantica, Tatsuya Morishima, Emily Mitchell, Maria Rosa Lidonnici, Fabienne Meier-Abt, Daniel Hayler, Laura Jardine, Abbie Curd, Muzlifah Haniffa, Giuliana Ferrari, Hitoshi Takizawa, Nicola K Wilson, Bertie Gottgens, Kourosh Saeb-Parsy, Mattia Frontini, and Elisa Laurenti, PhD. Unique molecular and functional features of extramedullary hematopoietic stem and progenitor cell reservoirs in humans. Blood, page blood.2021013450, January 2022.

[28] Anna S. E. Cuomo, Daniel D. Seaton, Davis J. McCarthy, Iker Martinez, Marc Jan Bonder, Jose Garcia-Bernardo, Shradha Amatya, Pedro Madrigal, Abigail Isaacson, Florian Buettner, Andrew Knights, Kedar Nath Natarajan, HipSci Consortium, Ludovic Vallier, John C. Marioni, Mariya Chhatriwala, and Oliver Stegle. Single-cell RNA-sequencing of differentiating iPS cells reveals dynamic genetic effects on gene expression. Nature Communications, 11(1):810, February 2020.

[29] Philip Ewels, Måns Magnusson, Sverker Lundin, and Max Käller. MultiQC: Summarize analysis results for multiple tools and samples in a single report. Bioinformatics, 32(19):3047–3048, October 2016.

[30] Christoph Hafemeister and Rahul Satija. Normalization and variance stabilization of single-cell RNA-seq data using regularized negative binomial regression. Genome Biology, 20(1):296, December 2019.

[31] Yuhan Hao, Stephanie Hao, Erica Andersen-Nissen, William M. Mauck, Shiwei Zheng, Andrew Butler, Maddie J. Lee, Aaron J. Wilk, Charlotte Darby, Michael Zager, Paul Hoffman, Marlon Stoeckius, Efthymia Papalexi, Eleni P. Mimitou, Jaison Jain, Avi Srivastava, Tim Stuart, Lamar M. Fleming, Bertrand Yeung, Angela J. Rogers, Juliana M. McElrath, Catherine A. Blish, Raphael Gottardo, Peter Smibert, and Rahul Satija. Integrated analysis of multimodal single-cell data. Cell, 184(13):3573–3587.e29, June 2021.

[32] Junyue Cao, Malte Spielmann, Xiaojie Qiu, Xingfan Huang, Daniel M. Ibrahim, Andrew J. Hill, Fan Zhang, Stefan Mundlos, Lena Christiansen, Frank J. Steemers, Cole Trapnell, and Jay Shendure. The single-cell transcriptional landscape of mammalian organogenesis. Nature, 566(7745):496–502, February 2019.

[33] Xiaojie Qiu, Qi Mao, Ying Tang, Li Wang, Raghav Chawla, Hannah A. Pliner, and Cole Trapnell. Reversed graph embedding resolves complex single-cell trajectories. Nature Methods, 14(10):979–982, October 2017.

[34] Ludo Waltman and Nees Jan van Eck. A smart local moving algorithm for large-scale modularity-based community detection. The European Physical Journal B, 86(11):471, November 2013.

[35] V. A. Traag, L. Waltman, and N. J. van Eck. From Louvain to Leiden: Guaranteeing well-connected communities. Scientific Reports, 9(1):5233, March 2019.

[36] Andrew D Yates, Premanand Achuthan, Wasiu Akanni, James Allen, Jamie Allen, Jorge Alvarez-Jarreta, M Ridwan Amode, Irina M Armean, Andrey G Azov, Ruth Bennett, Jyothish Bhai, Konstantinos Billis, Sanjay Boddu, José Carlos Marugán, Carla Cummins, Claire Davidson, Kamalkumar Dodiya, Reham Fatima, Astrid Gall, Carlos Garcia Giron, Laurent Gil, Tiago Grego, Leanne Haggerty, Erin Haskell, Thibaut Hourlier, Osagie G Izuogu, Sophie H Janacek, Thomas Juettemann, Mike Kay, Ilias Lavidas, Tuan Le, Diana Lemos, Jose Gonzalez Martinez, Thomas Maurel, Mark McDowall, Aoife McMahon, Shamika Mohanan, Benjamin Moore, Michael Nuhn, Denye N Oheh, Anne Parker, Andrew Parton, Mateus Patricio, Manoj Pandian Sakthivel, Ahamed Imran Abdul Salam, Bianca M Schmitt, Helen Schuilenburg, Dan Sheppard, Mira Sycheva, Marek Szuba, Kieron Taylor, Anja Thormann, Glen Threadgold, Alessandro Vullo, Brandon Walts, Andrea Winterbottom, Amonida Zadissa, Marc Chakiachvili, Bethany Flint, Adam Frankish, Sarah E Hunt, Garth IIsley, Myrto Kostadima, Nick Langridge, Jane E Loveland, Fergal J Martin, Joannella Morales, Jonathan M Mudge, Matthieu Muffato, Emily Perry, Magali Ruffier, Stephen J Trevanion, Fiona Cunningham, Kevin L Howe, Daniel R Zerbino, and Paul Flicek. Ensembl 2020. Nucleic Acids Research, 48(D1):D682–D688, January 2020.

[37] Yang Liao, Gordon K. Smyth, and Wei Shi. featureCounts: An efficient general purpose program for assigning sequence reads to genomic features. Bioinformatics, 30(7):923–930, April 2014.

[38] Mirjana Efremova, Miquel Vento-Tormo, Sarah A. Teichmann, and Roser Vento-Tormo. CellPhoneDB: Inferring cell–cell communication from combined expression of multisubunit ligand–receptor complexes. Nature Protocols, 15(4):1484–1506, April 2020.

[39] Kelly Street, Davide Risso, Russell B. Fletcher, Diya Das, John Ngai, Nir Yosef, Elizabeth Purdom, and Sandrine Dudoit. Slingshot: Cell lineage and pseudotime inference for singlecell transcriptomics. BMC Genomics, 19(1):477, June 2018.

[40] Tim Stuart, Avi Srivastava, Shaista Madad, Caleb A. Lareau, and Rahul Satija. Single-cell chromatin state analysis with Signac. Nature Methods, 18(11):1333–1341, November 2021.

[41] F. William Townes, Stephanie C. Hicks, Martin J. Aryee, and Rafael A. Irizarry. Feature selection and dimension reduction for single-cell RNA-Seq based on a multinomial model. Genome Biology, 20(1):295, December 2019.

[42] W. N. Venables, Brian D. Ripley, and W. N. Venables. Modern Applied Statistics with S. Statistics and Computing. Springer, New York, 4th ed edition, 2002.

[43] James Baglama and Lothar Reichel. Augmented Implicitly Restarted Lanczos Bidiagonalization Methods. SIAM Journal on Scientific Computing, 27(1):19–42, January 2005.

[44] Sunil Arya, David M. Mount, Nathan S. Netanyahu, Ruth Silverman, and Angela Y. Wu. An optimal algorithm for approximate nearest neighbor searching fixed dimensions. Journal of the ACM, 45(6):891–923, November 1998.

[45] Vincent D. Blondel, Jean-Loup Guillaume, Renaud Lambiotte, and Etienne Lefebvre. Fast unfolding of communities in large networks. Journal of Statistical Mechanics: Theory and Experiment, 2008(10):P10008, October 2008.

[46] Randolf Rotta and Andreas Noack. Multilevel local search algorithms for modularity clustering. ACM Journal of Experimental Algorithmics, 16:2.3:2.1–2.3:2.27, July 2011.

[47] Yasin S, enbabaoğlu, George Michailidis, and Jun Z. Li. Critical limitations of consensus clustering in class discovery. Scientific Reports, 4(1):6207, August 2014.

[48] Alexander J. Gates, Ian B. Wood, William P. Hetrick, and Yong-Yeol Ahn. Element-centric clustering comparison unifies overlaps and hierarchy. Scientific Reports, 9(1):8574, December 2019.

[49] Ruoxin Li and Gerald Quon. scBFA: Modeling detection patterns to mitigate technical noise in large-scale single-cell genomics data. Genome Biology, 20(1):193, December 2019.

[50] Philip Brennecke, Simon Anders, Jong Kyoung Kim, Aleksandra A. Kołodziejczyk, Xiuwei Zhang, Valentina Proserpio, Bianka Baying, Vladimir Benes, Sarah A. Teichmann, John C. Marioni, and Marcus G. Heisler. Accounting for technical noise in single-cell RNA-seq experiments. Nature Methods, 10(11):1093–1095, November 2013.

[51] D. Eppstein, M. S. Paterson, and F. F. Yao. On Nearest-Neighbor Graphs. Discrete & Computational Geometry, 17(3):263–282, April 1997.

[52] Jacob H. Levine, Erin F. Simonds, Sean C. Bendall, Kara L. Davis, El-ad D. Amir, Michelle D. Tadmor, Oren Litvin, Harris G. Fienberg, Astraea Jager, Eli R. Zunder, Rachel Finck, Amanda L. Gedman, Ina Radtke, James R. Downing, Dana Pe’er, and Garry P. Nolan. Data-Driven Phenotypic Dissection of AML Reveals Progenitor-like Cells that Correlate with Prognosis. Cell, 162(1):184–197, July 2015.

[53] V. A. Traag, P. Van Dooren, and Y. Nesterov. Narrow scope for resolution-limit-free community detection. Physical Review E, 84(1):016114, July 2011.

[54] Mirjana Efremova and Sarah A. Teichmann. Computational methods for single-cell omics across modalities. Nature Methods, 17(1):14–17, January 2020.

[55] Morgane Pierre-Jean, Jean-François Deleuze, Edith Le Floch, and Florence Mauger. Clustering and variable selection evaluation of 13 unsupervised methods for multi-omics data integration. Briefings in Bioinformatics, 21(6):2011–2030, December 2020.

[56] Lucy L Gao, Jacob Bien, and Daniela Witten. Are clusterings of multiple data views independent? Biostatistics, 21(4):692–708, October 2020.

[57] Tallulah S Andrews and Martin Hemberg. M3Drop: Dropout-based feature selection for scRNASeq. Bioinformatics, 35(16):2865–2867, August 2019.

[58] Etienne Becht, Leland McInnes, John Healy, Charles-Antoine Dutertre, Immanuel W. H. Kwok, Lai Guan Ng, Florent Ginhoux, and Evan W. Newell. Dimensionality reduction for visualizing single-cell data using UMAP. Nature Biotechnology, 37(1):38–44, January 2019.

[59] Brian Hie, Hyunghoon Cho, Benjamin DeMeo, Bryan Bryson, and Bonnie Berger. Geometric Sketching Compactly Summarizes the Single-Cell Transcriptomic Landscape. Cell Systems, 8(6):483–493.e7, June 2019.

[60] Huidong Chen, Caleb Lareau, Tommaso Andreani, Michael E. Vinyard, Sara P. Garcia, Kendell Clement, Miguel A. Andrade-Navarro, Jason D. Buenrostro, and Luca Pinello. Assessment of computational methods for the analysis of single-cell ATAC-seq data. Genome Biology, 20(1):241, December 2019.

[61] Ruben Chazarra-Gil, Stijn van Dongen, Vladimir Yu Kiselev, and Martin Hemberg. Flexible comparison of batch correction methods for single-cell RNA-seq using BatchBench. Nucleic Acids Research, (gkab004), February 2021.

